# NOT ALL ANTIBODIES HAVE BEEN CREATED EQUAL: Antibodies validated for routinely processed tissue seldom apply to frozen tissue sections

**DOI:** 10.1101/2020.05.10.087361

**Authors:** Maddalena M Bolognesi, Francesco Mascadri, Roberto Perego, Silvia Bombelli, Francesca M Bosisio, Giorgio Cattoretti

**Affiliations:** Pathology, Department of Medicine and Surgery, Università di Milano-Bicocca, Via Cadore 48, Monza 20900 (MI), Italy; Laboratory of Translational Cell and Tissue Research, KU Leuven, Herestraat 49, 3000 Leuven, Belgium; Dept. of Pathology, Ospedale San Gerardo, ASST-Monza, VIa Pergolesi 33, Monza 20900 (MI) IT

## Abstract

A customized context-dependent validation of antibodies for the prospective use, rather than a general stamp of validity, is required for reproducibility and data validity, besides the need of standardized reagents. In-situ antibody staining is a technique broadly used in experimental settings and human diagnostic practice. The first typically, but not exclusively uses lightly fixed material (cell smears, frozen sections), the second, routinely processed formalin-fixed, paraffin-embedded (FFPE) tissue. Differently from techniques based on tissue extraction, there is little awareness of the context-dependent constraints inherent with either type of in situ staining except that antigen masking associated with routine tissue processing limits the range of useful antibodies. We applied a panel of 126 antibodies validated for FFPE to lightly fixed (acetone, formalin) frozen sections and found that less than 30% performed conservatively with all fixations, 35% preferred one fixation over another, 13% gave non-specific staining, 23% did not stain at all. Individual antibody variegation of the paratope fitness for the differentially fixed antigen may be the cause. Re-validation of established antibody panels is required when applied to sections whose fixation and processing is different from the tissue where they have been initially validated.

## INTRODUCTION

Over the last fifteen years, awareness of the frequent variability and unreliability of data caused by the use of non-standardized, not validated antibodies (Abs) in various types of immunoassays such as Western Blot (WB), Immunocytochemistry (ICC) and Immunohistochemistry/immunofluorescence (IHC/IF) (1-4), has suggested common guidelines for the correct use of these reagents. An International Working Group for Antibody Validation (IWGAV) convened in 2016 and published a proposal to ensure reproducibility and validity when using antibodies for immunolabeling (5). Five validation principles and criteria were proposed, based on: stain reduction upon genetic ablation of the target (“genetic”), comparison with an antibody-independent method (“orthogonal”), comparison of two different Abs against the same target (“independent antibody”), comparison with a “tagged protein expression” and mass spectrometry analysis of the immunocaptured protein (“IMS”). The suitability of each validation method for several applications, including in situ staining methods (ICC and IHC) was reported as well (5). Guidelines for antibody reporting in publications were published (3, 6) around the same time.

These suggested guidelines apply to very diverse subjects in many science fields, including Pathology. Surgical Pathologists in particular are using IHC in an highly regulated diagnostic settings, where pre-analytical, analytical and reporting rules need to be strictly defined, because of the human health implications of these assays (7). Surgical Pathology Departments may routinely use >200 different primary antibodies and introduce about 20 new antibodies a year (7), however the focus of the antibody/assay validation is on technical reproducibility across the caseload rather than on the antibody validation as intended by the International Working Group mentioned above. The antibody specificity for the target is a given assumption, often provided by the company supplying the antibody or the assay. On the other hand, the type of tissue target in Surgical Pathology is overwhelmingly FFPE tissue sections, whose pre-analytical variables are also highly regulated (8). The modifications of the protein targets in FFPE material and the performance of antibodies in such conditions have been extensively published and are the basis for a successful use of FFPE tissue for diagnostic, prognostic and theragnostic purposes (9-11). Because of the robustness of tissue processing and immunoanalysis, archived FFPE human material and Surgical Pathology techniques are also the most abundant and diffuse tools for translational and basic science research.

The IWGAV recommendations are that “approaches for antibody validation must be carried out in an application- and context-specific manner.” (5), suggesting that antibody validation must be independent for extractive (e.g. WB, immunoprecipitation) versus in-situ technique (ICC, IHC) on the basis of the different modifications that the protein target undergoes in each procedure (see Supplementary Table 1 in (5)). Although the IWGAV makes a distinction between ICC and IHC, reports on the requirement for an independent antibody validation on either type of tissue type are scanty at best, while the underlying assumption in the vast majority of publications is that an antibody good for FFPE must be good for a frozen section or a cell preparation as well and vice versa. The only caveat is that antigen masking in FFPE tissue makes it harder to find antibodies working on both types of tissues. A handful of groups (12-14) addressed in a systematic fashion the difference in antibody performance in FFPE versus lightly fixed frozen tissue section, these latter believed at that time to be the “golden standard” (12) to assess distribution, intensity and, ultimately, validity of an antibody. In those publications, the focus was on the differential antigen retention with each fixation protocol, rather than questioning the antibody specificity, which was a given predefined value, also because coming from the Surgical Pathology practice.

We have previously investigated the epitope specificity and binding requirements of a large number of antibodies for human FFPE material by applying them to swine routinely fixed material (15). The results showed that about 50% did not react, despite complete identity of the epitope in both species. This may have occurred because of the fine requirement for some Abs of the conformation of the target side peptide chains not involved in the antigen binding (16) or because of non-identical juxtaposed adjacent proteins in the fixed complex. Of the Abs reacting with both species, we had evidence of species-specific negative effect of formalin fixation of BCL6, restricted to one antibody and not to others, while all did reacted on acetone-fixed frozen sections (15). In a separate investigation (11), we found that the epitopes for some antibodies were selectively destroyed by antigen retrieval (AR), possibly because of the conformational nature of the epitope.

These observations suggest that validated antibodies for FFPE sections may bear undisclosed paratope variations which may affect sensitivity, validity and usage when applied outside that context. Therefore we investigated a large panel of antibodies, validated on FFPE, on lightly fixed (acetone, formalin) frozen tissue sections. Antibodies for FFPE sections may detect linear epitopes (15) in a profoundly denatured protein mixture, thus we expected them to be working spectacularly on lightly fixed frozen material, but we found that this is quite often not the case.

## MATERIALS AND METHODS

### Human specimens

Fully anonymized human surgical specimen leftovers (pediatric tonsils, normal kidney, discarded serial sections from routinely processed formalin-fixed, paraffin-embedded (FFPE) material) were either snap-frozen in -80C chilled isopentane (Merck Life Science S.r.l.,Milano, Italy) or fixed overnight at RT in buffered 4% formaldehyde (Bio-Optica Milano Spa, Milano, Italy), processed through a graded ethanol gradient, then in xylene and embedded in molten paraffin for sectioning.

The study has been approved by the Institutional Review Board Comitato Etico Brianza, N. 3204, “High-dimensional single cell classification of pathology (HDSSCP)”, October 2019. Patients consent was obtained or waived according to article 89 of the EU general data protection regulation 2016/679 (GDPR) and decree N. 515, 12/19/2018 of the Italian Privacy Authority.

### Antibody validation

A list with all the primary antibodies validated for FFPE use can be found in in Supplementary Table 1. The type of validation of each reagent was listed essentially according to Edfors et al. and Uhlén et al. (5, 17) and modified for tissue staining. The following criteria were used:

- <u>Orthogonal</u>: an antibody uniquely identifying a cell in the tissue whose high-dimensional phenotype (18, 19) corresponds to a cell described by single cell RNA sequencing (i.e transcriptomics) and bearing the transcript corresponding to the target is considered validated.
- <u>Independent Antibody</u>: A) two antibodies directed against two separate epitopes of the same protein and having an identical staining pattern and/or co-localizing by double IF or B) an antibody uniquely identifying a cell in the tissue whose high-dimensional phenotype corresponds to a cell whose phenotype is defined by a multidimensional flow cytometry panel, are considered validated
- <u>Genetic</u>: an antibody whose staining or absence of staining corresponds to a genetically engineered ectopic expression or absence of the target is considered validated
- <u>Peptide Microarray</u>: an antibody recognizing its unique peptide target on peptide microarrays (20, 21) is considered validated
- <u>Cross-species</u>: an antibody whose reactivity is conserved across genomic sequence variations in another species (15) is considered validated

We added an additional criterion, “historic”, for antibodies whose widespread use in multiple applications (e.g. CD20) strongly suggests validity, despite lacking one of the criteria listed above.

Evidence of validation not present in the published literature, either peer-reviewed papers or documentation from producers, was produced in house. Some antibodies were validated according to multiple criteria. Validated, FFPE-proof antibodies directed against the same protein can be used exchangeably on routinely processed sections, besides obvious variations in host species or isotype (22).

Representative iconography of the FFPE staining for each antibody can be viewed in published papers ((19, 23)), in (24) or at the Human Protein Atlas website (https://www.proteinatlas.org/).

### Frozen tissue fixation

Frozen tissue blocks were sectioned at 5 µm thickness in a Leica CM1850 cryostat (Leica Microsystems GmbH, Wetzlar, Germany) at -20C. Sections were collected on positively charged slides (Menzel-Glaser Superfrost Plus; Bio-Optica Milano Spa).

Two fixation methods were chosen: in one, sections on slides were lifted from the cryostat, placed temporarily in a slide rack, immersed in acetone (Merck) at +4C for 5 min, allowed the acetone to evaporate, stored at RT overnight in a damp-proof box; subsequently, the sections were either used the same day or stored wrapped in plastic foil at -20C for later use.

In the second method, slides with the frozen section attached were immediately immersed in 4% buffered formaldehyde (FA), fixed at RT for 18h (overnight), then washed in Tris-containing buffer for formaldehyde quenching (25). These sections were stored at +4C in 50 mM Tris HCl buffer pH 7.5, containing 0.01% Tween-20 (Merck) and 100 mM sucrose (TBS-Ts).

Previous published experiments (11) showed that fixation in excess of 30 min is required for tissue stabilization and epitope rescue upon AR and 48h or 72h fixation do not add additional masking, compared to overnight fixation, therefore we chose this latter fixation time for all the experiments.

Frozen sections were dipped for 5 min in 2.5% horse serum in TBS (Vector Laboratories, Burlingame, CA, USA) before use for Fc receptor blocking.

### Antigen retrieval

Antigen retrieval (AR) was performed placing the sections in a 800 ml glass container filled with the retrieval solutions (10 mM EDTA in Tris-buffer pH 8, Merck), irradiate in a household microwave oven at full speed for 8 min, followed by intermittent electromagnetic radiation to maintain constant boiling for the set time, and cooling the sections to about 50 °C before use.

We found that 20 min of AR on FA-fixed frozen sections did not produce any additional retrieval compared to 1 min or 5 min (not shown), thus a protocol comprising the 8 min preheating and 1 min AR was used throughout.

### Immunolabeling

Sections were processed for indirect immunohistochemistry or immunofluorescent labeling as previously described in detail (19, 23).

Briefly, the sections were incubated for 30 min with optimally diluted primary antibodies in combination of up to four (22), washed and counterstained with specific distinct fluorochrome-tagged secondary antibodies (23). The slides, counterstained with DAPI and mounted, were scanned on an S60 Hamamatsu scanner (Nikon, Italia) at 20x magnification (23). Optimal exposure time for each antibody-fluorochrome combination was set in advance on FFPE and validated over multiple samples in a multiplex experiment on routinely processed material ((18) and not shown). Quadruple simultaneous IF staining guarantees internal control for effective staining. Dubious results were confirmed in single stain IHC. The vast majority of the tests were done on tonsil sections, which constitute the optimal positive control for a panel in which anti-leukocyte antibodies are overrepresented. Acetone-fixed, formalin-fixed and formalin-fixed AR treated frozen sections were all processed simultaneously.

Results were scored semi-quantitatively (neg, +/-, +, ++, +++) and qualitative staining features (diffuse, non-specific staining, etc.) were annotated.

### Preparation of immunofluorescent images for image analysis

Single .ndpi images for each case were saved as .tiff files and autofluorescence (AF) was subtracted (23). Grayscale images were inverted, brightness and contrast adjusted with the default ImageJ function (26), simultaneously processing a stack of three experimental sets (acetone, FA, FA>AR) before producing individual images.

## RESULTS

The overwhelming majority of antibodies tested were selected for being reactive with AR-treated FFPE sections (118/126), yet 8 (6.3%) selectively failed to stain acetone fixed sections and additional 8 had minimal reactivity compared to FA fixed sections.

Fourteen, including these latter eight, required formalin fixation to be detected (11.1%) (Table 1, Fig. 1 and labelled “FA dependent” in Supplementary Table 1) and AR was able to enhance the staining in only three of them.

**Table 1.** Summary table of the staining results, grouped by category.

| Reactivity | Acetone | FA | FA > AR | N. Abs | % | Note |
| --- | --- | --- | --- | --- | --- | --- |
| conserved | pos | pos | pos | 35 | 27.8% | Reactivity preserved on all fixations |
| FA blocked | pos | neg | pos | 25 | 19.8% | Inactivated by FA fixation (rescued by AR) |
| FA dependent | neg | pos | pos | 14 | 11.1% | Requires FA fixation |
| conformational | pos | pos | neg | 17 | 13.5% | Heat-induced appearance/disappearance |
|  | neg | neg | pos |  |  |  |
| misc | pos/neg | neg/pos | neg | 6 | 4.8% | Mostly Pos in Acetone fixed only |
| neg | neg | neg | neg | 29 | 23.0% | No reactivity |
| Total |  |  |  | 126 |  |  |

**Figure 1.**
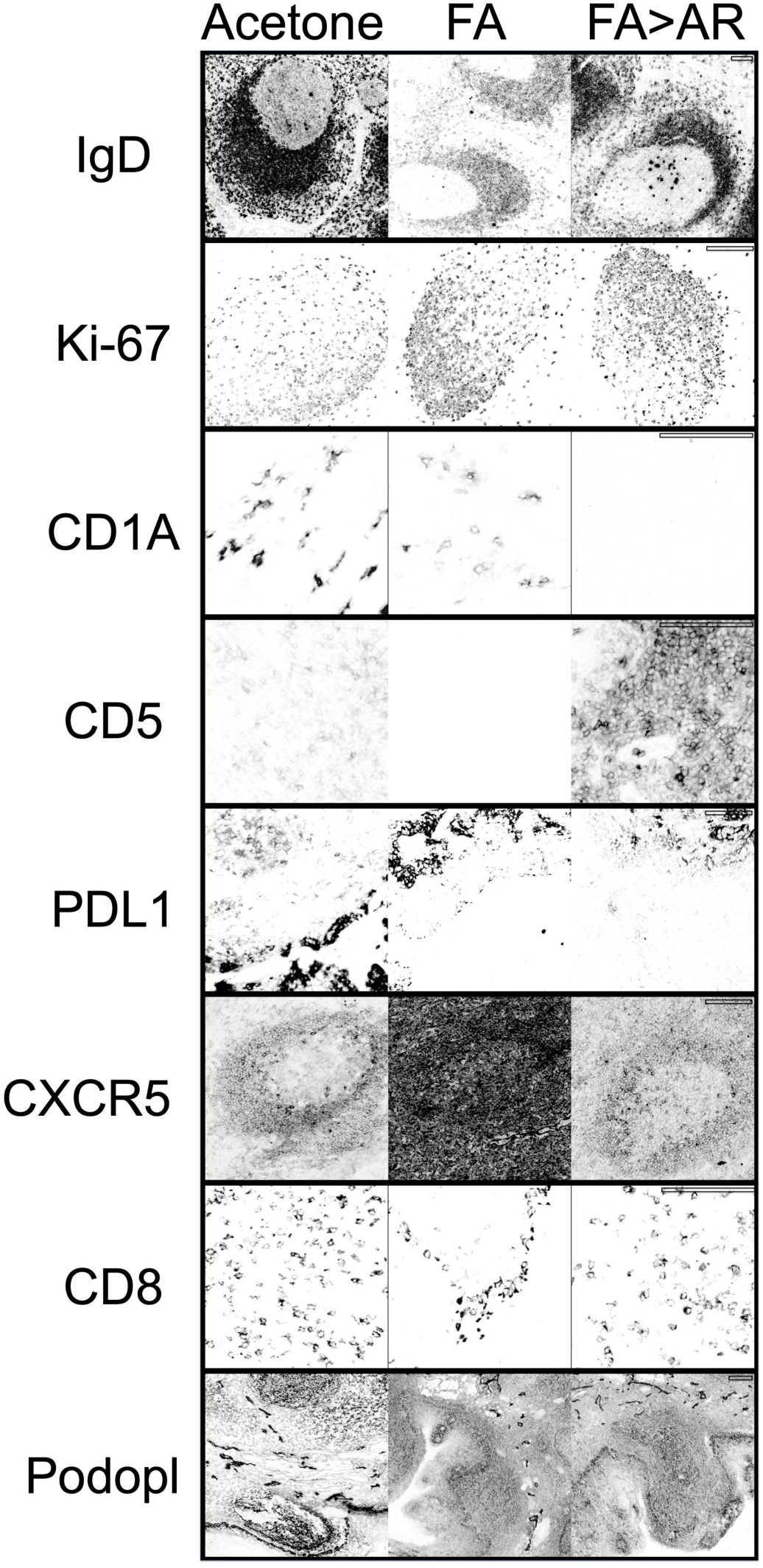
Examples of antibody staining on lightly-fixed frozen tonsil tissue. Representative examples of stainings obtained on acetone, formalin fixation (FA), and FA fixation followed by AR (FA>AR), left, middle and right columns respectively. From top to bottom: IgD (goat poly) and Ki-67 (UMAB107) are conserved in intensity across the fixations, the former reduced after FA fixation. CD1A (O10) staining is lost upon heating (AR). CD5 (CD5/54/F6) staining is lost upon FA fixation and regained after AR. PDL1: macrophage and epithelial cell staining for PDL1 (cocktail of rabbit MAbs) is maximal in Acetone-fixed section. Macrophage stain is all but gone in FA fixed, but partially retrieved after AR. Epithelial staining is preserved in FA and a basal layer staining is added, this latter lost upon AR. CXCR5: formalin fixation causes a diffuse non-specific staining for CXCR5 (#51505). Lymphocyte CD8 (C8/144B) staining is maintained and specific, except upon FA fixation, where is greatly reduced, while a non specific strong staining of the basal epithelial layer is produced. Podoplanin (NZ-1.2) produces a bright staining of the basal epithelium, endothelial cells and follicular dendritic cells in acetone-fixed cells. Upon FA fixation, endothelial cells remain strong, while the other structures are quite weaker. Scale bar = 50 µm.

FA fixation did reduce epitope detectability in a quarter of the cases and abolished the reactivity in as many as 24 cases (19.8%) (“FA blocked” in Table 1 Fig. 1 and Supplementary Table 1); in all these instances, AR rescued the optimal reactivity.

The epitope composition of the “FA blocked” antibodies was available for 5 Abs only and short enough to be informative in one (CD3-12; ERPPPVPNPDYEPC). The epitope contained no lysine, the topmost aminoacid bound by formaldehyde, yet was sensitive to FA fixation on frozen sections.

A group of 17 abs (13.5%) had immunoreactivity either selectively elicited by AR treatment (12 Abs) or abolished by the heat treatment. We defined those as “conformational” because of a possible disruption (or creation) of an epitope by altering the conformation of one or more protein loops generating the epitope (Table 1, Fig. 1 and Supplementary Table 1)

A group of 6 Abs (4.8%) we named “miscellaneous” (Table 1, and Supplementary Table 1) contained Abs behaving differently than the previous groups and were composed mostly of Abs reacting with acetone-fixed sections only. One antibody (GZMB) labelled selectively FA-fixed frozen sections.

Twenty nine Abs (22.8%) did not stain any type of lightly fixed sections.

The remaining Abs (35/126; 27.8%) had conserved reactivity across the three preparations: acetone fix, FA fix, FA>AR (“conserved”; Table 1, Fig. 1 and Supplementary Table 1).

Whenever the protein distribution was shared with cells of diverse origin (e.g. podoplanin on endothelium, basal epithelium and follicular dendritic cells), differential response to the fixation protocols was noted according to the cell type. As an example, only the endothelial staining of Podoplanin (Fig. 1) conserved an equivalent intensity across the three fixation protocols. For some protein, a loss of the target from sections in acetone fixed tissue was noticed (e.g. LYZ, S100AB)

Spurious reactivity, either due to non-specific, background staining or to discrete unrelated targets (Fig. 1) was detected in 17 instances (13.4%), 10 of them (7.8%) in FA fixed sections.

In 19 instances, the same protein was tested with multiple independent antibodies which showed in all instances but two individual variations in staining pattern (Supplementary Table 1).

Because of the antibody panel choice, biased toward leukocyte antigens, most Abs did not react with swine FFPE tissue (15), thus no epitope variation information could be obtained.

## DISCUSSION

A minority of antibodies validated for the use on FFPE material apply for the use on frozen sections, formalin- or acetone fixed. The most likely reason for such unexpected results may lie in the subtle requirement of the paratope for an unique epitope binding environment, composed not just of the epitope, but also of the neighboring residues on the protein sequence (16) or of adjacent unrelated proteins and their post-translational modifications (27). Such epitope microenvironment is affected by the fixation method and, when applied, by further denaturation provided by high heath.

So far, the burden of the proof has been put on the antigen and its modification during fixation. Our investigation however started from anti-peptide antibodies validated on the new “gold standard”, FFPE material (12), went back to the frozen tissue perceived as containing native proteins, and has unveiled how context-dependent the use of Abs for in situ detection can be.

Differently from previous published work (12-14), the use of multiple antibodies for the same target (35% of the panel) has disclosed a remarkably heterogeneous staining ability within the same target across the experimental conditions, suggesting that the source of variance is the paratope variegation, rather than epitope.

It has also shown evidence of the persistence of the antigen in the tissue with that particular fixation condition, indirectly re-validating the use of acetone-fixed or formalin-fixed frozen sections as “alternative gold standards” for antigen detection in tissue. Only a couple of antigens may have been lost from acetone-fixed sections, as previously published (12, 13).

The recommendation to validate and use antibodies limited to the technique employed (Western Blot, immunoprecipitation, immunohistochemistry, etc.) (5, 17) may now be made in a more detailed fashion also for immunological in situ techniques, distinguishing frozen sections from FFPE material, and within frozen sections, whether acetone of FA is used and if AR is required. Particularly worrisome for staining reliability across fixations is the presence, albeit small, of unrelated discrete reactivity (e.g. CD8 on epithelia; this report and (28, 29)) or the variegated effect of treatment on epitopes shared across diverse cell types (e.g. Podoplanin, PDL1) (Fig. 1).

The major bias of this study is the antibody choice, which was guided by the validity on routinely processed tissue sections, being only 9 of the selected Abs not reactive on FFPE. Another bias is the abundance of mouse non-IgG1, non-rabbit antibodies (42%), due to the multiplexing staining strategy (22), selection which however may have highlighted the problem by diversifying, besides the immunizing epitope, the host view of the immunogen.

Collectively, these data suggest that antibody binding may be very dependent on tissue fixation and processing in a subtle and unexpected fashion, affecting binding and specificity for the target.

## ACKNOWLEDGEMENTS

This work has been supported by the Departmental University of Milano-Bicocca funds and by a Fondazione Cariplo grant (2017-0577) to Prof. Roberto Perego. MMB was funded by a Cariplo grant (2017-0577) and is a PhD student in the DIMET PhD Program call XXXV of the Department of Medicine and Surgery of the University of Milano-Bicocca since November 2019. We wish to thank Carlo Parravicini for suggestions, Lorella Riva and Loredana Tusa for expert help in histopathology.

## AUTHOR’S CONTRIBUTIONS

GC and MMB equally designed the experiments.

MMB devised the image analysis algorithms, performed visual and digital image analysis as well as bioinformatic evaluation.

FM and SB performed immunostaining experiments. FMB and RP provided essential reagents.

MMB, FMB and GC wrote the manuscript.

All authors have read and approved the final manuscript.

## SUPPLEMENTARY MATERIAL

**SUPPLEMENTARY TABLE 1.**
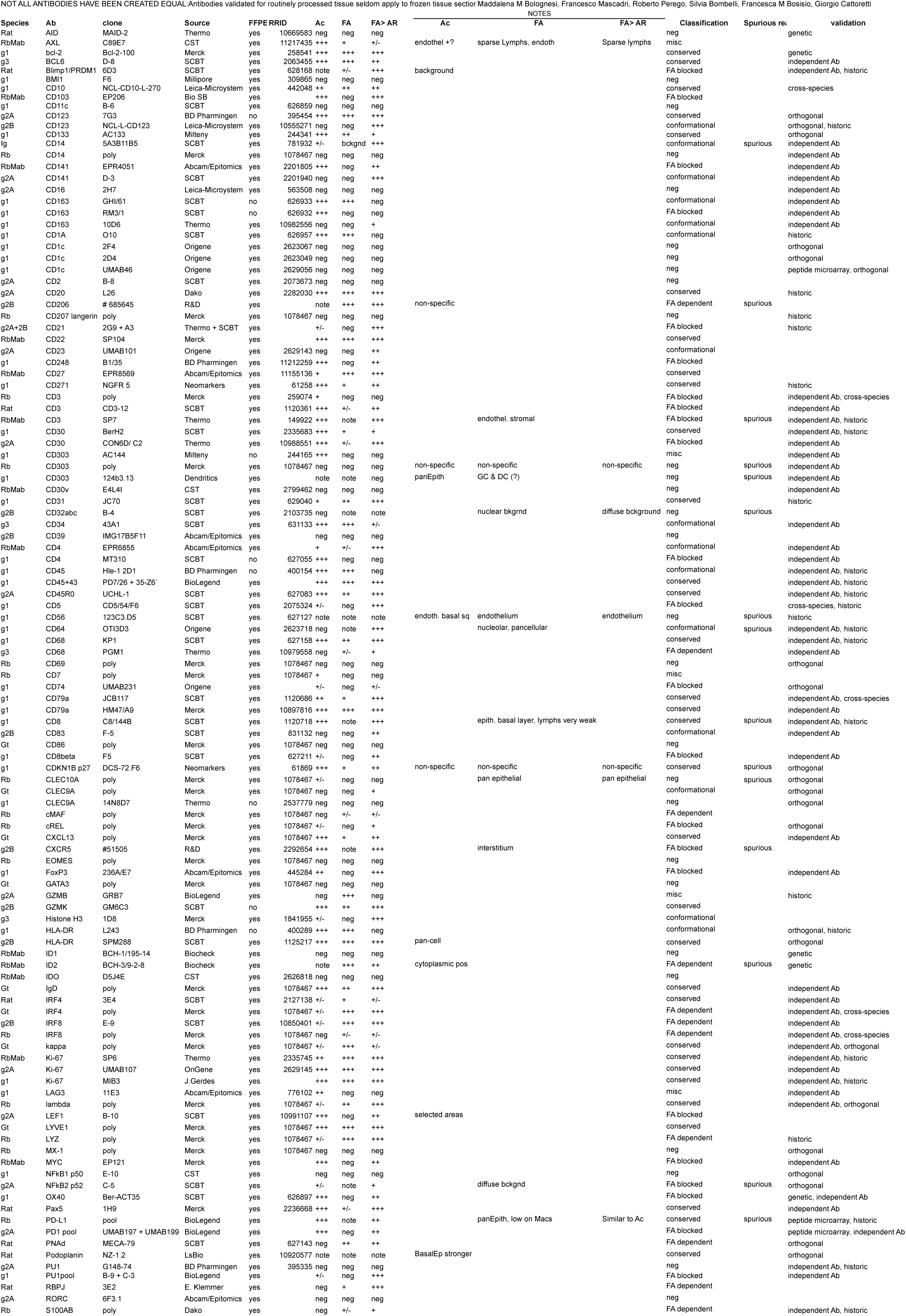
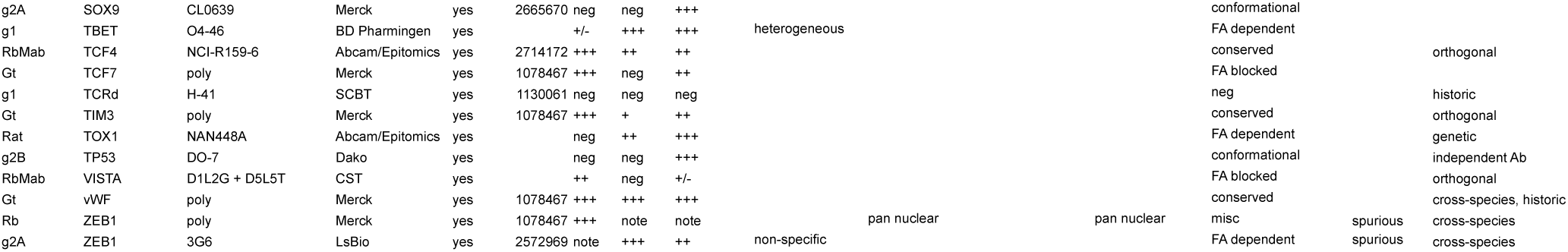
List of primary antibodies. Abbreviations: Species: Rb= rabbit; RbMab= rabbit monoclonal ab; g1, g2a, g2b, g3= mouse isotypes. FFPE: yes= working on routinely processed material. RRID: Resources Identification Portal identifier (https://scicrunch.org/resources) Classification: conformational= conformational epitope; conserved= staining conserved over fixation types; FA blocked= reactivity blocked by FA fixation; FA dependent= staining depends on FA fixation; negative= no staining; misc= miscellaneous behaviour, mostly Acetone only, not comprised in the previous classification. Spurious= spurious, non-specific reactivity present. Validation: type of antibody validation as described in M&M.

